# RNAGym: Large-scale Benchmarks for RNA Fitness and Structure Prediction

**DOI:** 10.1101/2025.06.16.660049

**Authors:** Rohit Arora, Murphy Angelo, Christian Andrew Choe, Courtney A. Shearer, Aaron W. Kollasch, Fiona Qu, Ruben Weitzman, Artem Gazizov, Sarah Gurev, Erik Xie, Debora S. Marks, Pascal Notin

**Author notes:** Equal contributions (Rohit Arora, Murphy Angelo, Christian Andrew Choe, Courtney A. Shearer). Senior authorship (Debora S. Marks, Pascal Notin).

## Abstract

Understanding RNA structure and predicting the functional consequences of mutations are fundamental challenges in computational biology with broad implications for therapeutic development and synthetic biology. Current evaluation of machine learning-based RNA models suffers from disparate experimental datasets and inconsistent performance assessments across different RNA families. To address these challenges, we introduce RNAGym, a large-scale benchmarking framework specifically designed for three core tasks–RNA fitness, secondary structure, and tertiary structure prediction. The framework integrates extensive datasets, including 70 standardized deep mutational scanning assays covering over a million mutations across diverse RNA types; 901k chemical-mapping reactivity profiles for secondary structure; and 215 diverse tertiary structures curated from the PDB. RNAGym is designed to facilitate a systematic comparison of RNA models, offering an essential resource to enhance the understanding and development of these models.

## 1 Introduction

RNA, once considered a passive intermediate between DNA and protein, is now recognized as a dynamic and crucial agent in cellular process regulation. The complexity of RNA lies in its structure — the base-pairing patterns that form the backbone as well as its two- and three-dimensional architecture— and in its functional versatility, which ranges from catalytic activities to gene regulation. Predicting RNA structure and assessing the functional impact of sequence variations, ie. RNA fitness, are key challenges in computational biology and machine learning. These interrelated tasks are critical for advancing our understanding of RNA biology and its applications in fields such as drug discovery and synthetic biology.

The prediction of RNA structure remains a significant challenge. While computational methods have made substantial progress, they still face considerable hurdles, especially for larger RNAs (>100 nt) with complex features like multi-branched loops and pseudoknots. Experimental methods like nuclear magnetic resonance, Cryo-EM, and X-ray crystallography can determine RNA 3D structures, but they face technical limitations when applied to RNA, resulting in RNA structures comprising less than 1% of entries in the Protein Data Bank (PDB) [Burley et al., 2017]. This scarcity of experimental data further complicates the development and validation of computational prediction methods.

An equally critical task is predicting RNA fitness – the functional capacity of RNA sequences when subjected to mutations. Understanding the impact of these mutations on RNA function is crucial for advancing our knowledge of RNA evolution and its role in cellular processes. This task is also vital for the development of RNA-based therapeutics and the expansion of synthetic biology applications, such as designing riboswitches for gene regulation or engineering RNA sensors for metabolite detection. Despite its importance, accurately predicting the functional consequences of RNA mutations, especially from sequence data alone, remains a significant challenge in the field. Both structure and fitness prediction can benefit from evolutionary information. For structure prediction, approaches such as maximum entropy models leverage sequence co-variation to infer evolutionary constraints [Weinreb et al., 2016, Hopf et al., 2017, Frazer et al., 2021]. Similarly, fitness prediction methods can utilize evolutionary data to identify functionally crucial sequence features. However, robust methodologies for integrating this information and accurately predicting both structure and fitness, particularly in zero-shot scenarios, remain elusive.

To address these challenges and support progress in the field, we present RNAGym, a comprehensive benchmarking framework designed to evaluate and compare computational methods for RNA structure and fitness prediction. We curate a large collection of deep mutational scanning assays, chemical mapping datasets for secondary structure prediction, and tertiary structures from the PDB. By providing a systematic way to evaluate model performance across various RNA types and prediction tasks, RNAGym can help identify strengths and weaknesses of current methods, guide the development of more accurate algorithms, and ultimately contribute to advancing our understanding of RNA biology and its applications in areas such as personalized medicine, RNA-based drug design, and engineered RNA devices for synthetic biology.

## 2 Related Work and Background

### 2.1 Prior RNA benchmarks

Existing RNA benchmarks for noncoding variant effect prediction have been limited and fragmented, primarily focusing on testing individual models rather than serving as comprehensive benchmarking platforms. For instance, the RfamGen model was evaluated using five assays, including datasets on ribozymes and tRNAs [Sumi et al., 2024]. Similarly, the Evo model was assessed using nine assays, incorporating ncRNA mutational scanning datasets [Nguyen et al., 2024]. Both studies relied on overlapping but distinct datasets to evaluate their models, making it difficult to compare performance metrics between studies directly. These small benchmark sets restrict the ability to generalize findings and were primarily used to test the respective models’ performance, rather than providing a broad and standardized benchmarking framework spanning the diversity of RNA types. This limited scope stands in stark contrast to the field of protein research, where platforms like ProteinGym have been established to offer extensive and standardized benchmarking datasets [Notin et al., 2023]. RNAGym addresses this gap by introducing a comprehensive benchmarking platform for RNA variant effect prediction that offers more than four times the number of assays compared to previous efforts, across a broader array of RNA classes. Previous RNA benchmarks include BEACON Ren et al. [2024], which covers 13 diverse tasks spanning secondary structure, functional genomics, and RNA engineering, and RNABench Runge et al. [2024], which provides six benchmarks centered on RNA structure prediction and design. In contrast, RNAGym specifically targets RNA fitness (variant effect) prediction and RNA folding tasks across both secondary and tertiary structures. With respect to 3D RNA structure prediction, several competitive benchmarks have been developed, including the Critical Assessment of Structure Prediction (CASP) and RNA-Puzzles. CASP15 [Kryshtafovych et al., 2023], the latest iteration of CASP, introduced a dedicated category for RNA structure prediction, reflecting the growing recognition of RNA’s importance and the need for accurate computational models. RNA-Puzzles [Cruz et al., 2012], on the other hand, is a community-driven initiative that presents real-world challenges to participants, who submit their models to be evaluated against experimentally determined RNA structures. Notably, only a few RNA molecules are evaluated at CASP and through RNA-Puzzles, limiting the ability of these high quality datasets to act as benchmarking standards. Train-test splits are commonly used to evaluate 2D RNA structure prediction models [Penić et al., 2024, Chen et al., 2022]. Intra-family splits involve training models on RNA sequences from the same family, with sequences from these families appearing in both the training and test sets [Singha et al., 2019]. This approach tests a model’s ability to learn and predict structures within known families. In contrast, inter-family splits ensure that sequences from the same RNA family in the test set are excluded from the training set [Penić et al., 2024]. This method assesses whether a model can generalize to entirely new RNA families that were not included in the training data. While existing benchmarks offer valuable insights, they often lack the scale and diversity to comprehensively evaluate model performance across various RNA types and structures. Zero-shot benchmarks for structure prediction models are crucial, as they avoid biases inherent in supervised approaches and potential overfitting, thus providing a more robust assessment of true generalization capabilities.

### 2.2 Background: the diversity of RNA molecules

RNA molecules exhibit a remarkable range of structures and functions, highlighting their essential role in both the fundamental processes of biology and their growing utility in medical and biotechnological applications. **mRNAs (Messenger RNAs)** act as the intermediary transcript that carry genetic information from DNA to the ribosome, where they serve as a template for protein synthesis. The sequence of mRNA dictates the amino acid sequence in a protein, thereby directly influencing gene expression and regulatory mechanisms. While alterations in splicing impact how exons are joined together to form the mature mRNA transcript, missense changes directly alter the codon sequence itself, resulting in the incorporation of a different amino acid in the final protein product. **tRNAs (Transfer RNAs)** play a crucial role in translation, the process of protein synthesis in the ribosome. Each tRNA molecule transports a specific amino acid to the ribosome; its anticodon loop pairs with the corresponding codon in the mRNA, ensuring that the correct amino acid is incorporated into the growing protein chain. **Ribozymes** are catalytic RNA molecules that perform specific biochemical reactions akin to protein enzymes. These include critical activities such as RNA splicing during gene expression, where ribozymes help in the excision of introns from a pre-mRNA. **Aptamers** are short, single-stranded RNA molecules designed to bind with high specificity and affinity to certain targets, including proteins, small molecules, and various cellular components.

## 3 RNAGym benchmarks

### 3.1 Overview

RNAGym is a comprehensive benchmark suite advancing the development and analysis of machine learning RNA models. It comprises three integrated layers: datasets, models, and analytics (Fig. 1), supporting three core RNA tasks:

- **Fitness prediction:** Prediction of RNA functionality across diverse RNA types, leveraging a broad set of deep mutational scanning assays.
- **Secondary Structure prediction:** Prediction of RNA secondary structure, focusing on identifying nucleotide contacts, which is crucial for understanding RNA function.
- **Tertiary Structure prediction:** Prediction of RNA tertiary structure, extending beyond secondary contacts to evaluate critical higher-order interactions.

**Figure 1:**
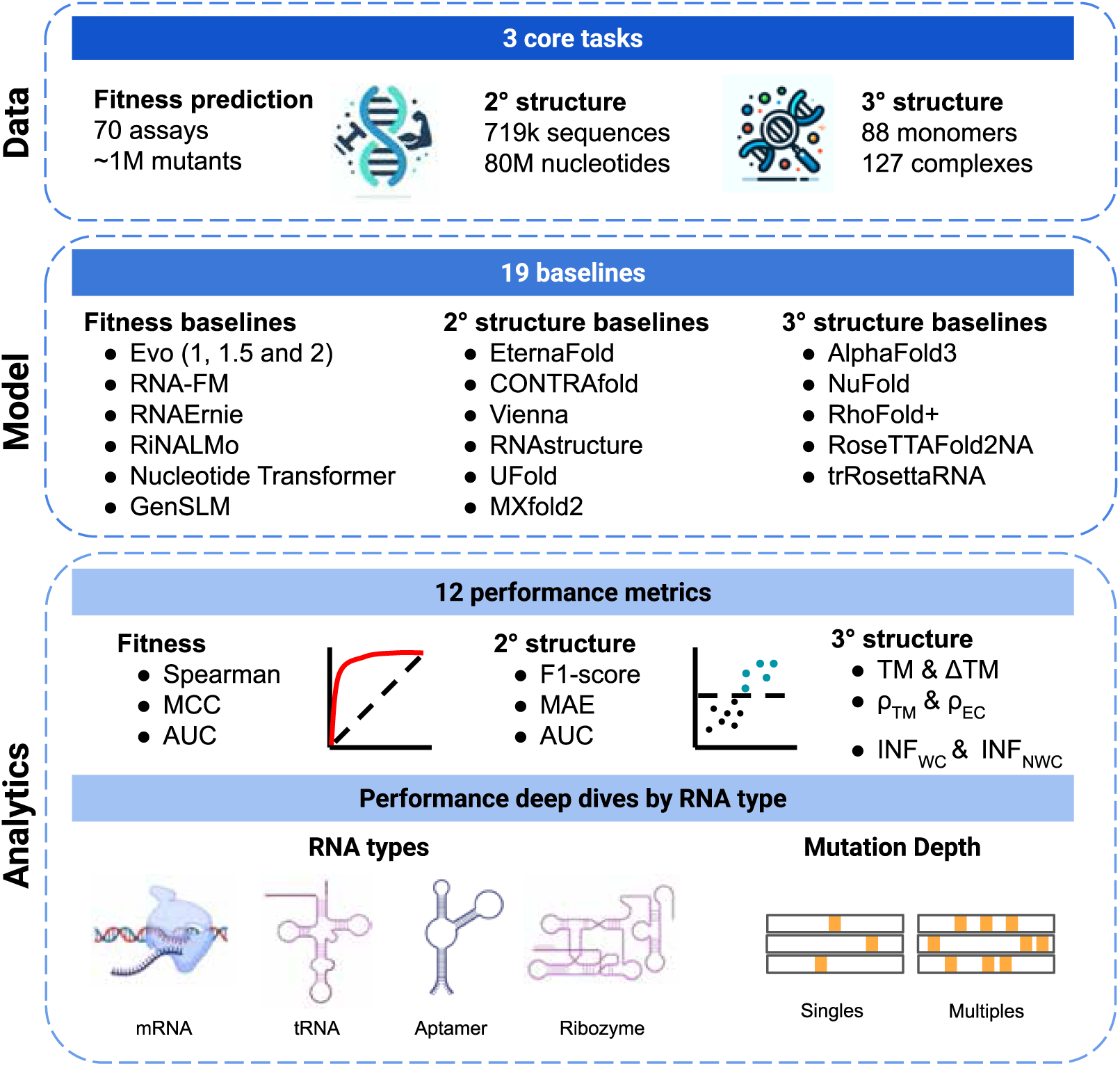
RNAGym benchmarks. RNAGym is a comprehensive RNA analysis framework designed specifically for fitness and structure prediction tasks. It evaluates the performance of diverse baselines across these tasks, and offers in-depth assessments by RNA type and mutation depth.

These tasks, all evaluated in a **zero-shot setting**, challenge models to generalize across varied RNA contexts without task-specific fine-tuning. Our data layer includes curated datasets that are specifically structured for these three tasks. These datasets are enriched with detailed annotations for a variety of RNA types and are classified by mutation depth, enhancing the granularity of the data available for analysis. Across all tasks, RNAGym integrates a diverse array of 19 predictive models, each tailored to address the nuances of the specific tasks at hand—whether predicting RNA fitness or determining RNA structure. The analytics layer of RNAGym is designed to provide a deep and comprehensive evaluation of model performance. It utilizes eleven distinct performance metrics to assess the effectiveness of each model in a clear and quantifiable manner. Further, the framework allows for detailed exploration of model performance across different RNA types and mutation depths, with the goal of understanding model strengths and limitations in varied biological contexts.

### 3.2 Datasets

#### Fitness prediction

For noncoding RNA studies, we conducted a broad PubMed search yielding over 11k results, which we pre-screened using a LLM, and then manually reviewed with specific inclusion/exclusion criteria as detailed in Appendix B. Coding mRNA datasets were sourced directly from ProteinGym when nucleotide-level deep mutational scanning information was available, aiming for a balanced set across taxa and assayed function types. RNAGym includes 70 Deep Mutational Scanning assays containing 1,117,995 variants measuring fitness. Beyond the sheer size of RNAGym, assays span a broad range of experimental contexts from in vivo cellular fitness measurements to in vitro biochemical assays across various tRNA, aptamers, ribozymes, and both coding and splicing disrupting mRNAs (Table 1). This effort represents a *fourfold increase* in size over the largest prior RNA benchmarks for non coding fitness prediction Brixi et al. [2025]. Unlike previous efforts, these assays are integrated into a standardized, reusable public resource, making RNAGym a more accessible and broadly applicable tool for RNA fitness prediction. Our assays all follow a standard format (see Appendix A).

**Table 1:** Fitness prediction assays. Overview of the diverse set of DMS assays in RNAGym, showing the number of assays and number of single and multiple mutants per RNA type.

| RNA Type | Description | # Assays | # Singles | # Multiples |
| --- | --- | --- | --- | --- |
| Aptamer | Target binding ability | 2 | 0.4k | 2.6k |
| Ribozyme | Cleavage and splicing | 26 | 4.1k | 755k |
| Transfer RNA (tRNA) | Stability and growth | 3 | 0.4k | 95k |
| Messenger RNA (mRNA) | Splicing ability | 2 | 0.3k | 5.4k |
| Messenger RNA (mRNA) | Coding mRNA fitness | 37 | 40k | 215k |
| Total |  | 70 | 46k | 1,072k |

#### 2° structure prediction

In preparing the benchmark for our research paper, we utilized the dataset from the Stanford Ribonanza Challenge, which contains chemical mapping data for many RNA sequences. Chemical mapping is a way to chemically probe an RNA structure by reacting the RNA with various reagents, commonly DMS (dimethyl sulfate) and 2A3 (2-aminopyridine-3-carboxylic acid imidazolide). Once an RNA is reacted with the chemical mapping reagent, the reactivity of each nucleotide can be read by reverse transcription and high-throughput sequencing. The reactivity profile gives a one dimensional view of the RNA structure where high reactivity is associated with single-stranded regions (unpaired RNA) and low reactivity is associated with double-stranded regions (paired RNA) [Spitale and Incarnato, 2023]. This type of data is invaluable for validating computational models of RNA secondary structure prediction, as it offers direct evidence of the RNA’s physical structure (Table 2). For completeness, we included data on ligand binding, cotranscriptional folding, and RNA degradation. We compiled the data from RMDB [Cordero et al., 2012] and filtered for the sequences with signal-to-noise ratio ≥ 1.0. For our analysis we focused on only in vitro chemical mapping data involving DMS and 2A3 for ease of interpretation. This resulted in a high quality dataset with 901*k* reactivity profiles for 583*k* unique sequences and 100*M* nucleotides (Table A10). The test dataset provides a score for each nucleotide for all RNA sequences (1 row per nucleotide), reflecting the propensity of that nucleotide to be single-stranded in the RNA structure.

**Table 2:** 2° structure dataset. The secondary structure dataset consists of RNA reactivity data compiled from RMDB. Standard RNA refers to the experiment done in vitro.

| Dataset Type | Description | # Sequences | # Filtered Sequences |
| --- | --- | --- | --- |
| 2A3 | Standard RNA | 2,136k | 471k |
| DMS | Standard RNA | 2,148k | 411k |
| Other modifier | Standard RNA | 70k | 47k |
| Ligand binding | RNA with ligands | 292 | 292 |
| In vivo | RNA is folded in vivo | 30 | 29 |
| Cotranscriptional | RNA is folded cotranscriptionally | 3k | 3k |
| Degradation | Accelerated RNA degradation | 18k | 16k |
| Total |  | 4,375k | 947k |

#### 3° structure prediction

RNAGym also introduces a 3D RNA structure dataset that emphasizes evaluation across a wide range of lengths, functional classes, and degree of structure and sequence homology to training data (Table 3). Starting with all 21,000+ RNA structures in the PDB, we filtered for RNAs published after January 12, 2023, ensuring no overlap with any baseline’s training set. The remaining RNAs were annotated with **TM_train_**and **ID_train_**, representing the maximum structure and sequence homology, respectively, of each evaluation candidate to any RNA published prior to the aforementioned date cutoff. Finally, the top three structures of each RNA family—ranked by TM_train_, ID_train_, length (up to 2,000 nucleotides), and resolution (up to 5.0Å)—were included in the overall dataset. These metrics were paired with unsupervised evolutionary information from EVCouplings to contextualize model performance. This framework allows us to assess not only how well models predict a variety of RNA tertiary structures but also how they generalize to novel folds and how well they extract coevolutionary signals.

**Table 3:** 3° structure dataset. RNAGym features diverse 3D RNA monomers and complexes. Rfams:= unique top-matching RNA families (E-value ≤ 1). Rfam signatures capture structural diversity of unique top-matching Rfam footprints (E-value ≤ 10). Monomers := chains with <33% residues bound to another polymer (within 4.5Å). Complexes := all others.

| Dataset | # Chains | # Rfams | # Rfam signatures | Avg. resolution | Avg. length |
| --- | --- | --- | --- | --- | --- |
| Monomers | 88 | 37 | 60 | $2.94\text{\AA}$ | 130 nt |
| Complexes | 127 | 32 | 74 | $2.98\text{\AA}$ | 52 nt |
| RNAGym (total) | 215 | 66 | 127 | $2.96\text{\AA}$ | 84 nt |

### 3.3 Baselines

We benchmarked a diverse set of RNA models across the three prediction tasks. For **fitness prediction**, we evaluated both large-scale foundation models trained on genomic data, including EVO 1 (7B parameters) [Nguyen et al., 2024], EVO 1.5 [Merchant et al., 2024], EVO 2 [Brixi et al., 2025], and Nucleotide Transformer [Dalla-Torre et al., 2023], alongside smaller specialized models focused on non-coding RNAs such as RiNALMo (650M parameters) [Penić et al., 2024], RNA-FM [Chen et al., 2022], GenSLM [Zvyagin et al., 2023], and RNAernie [Wang et al., 2024a]. For **2° structure prediction**, we included both classical thermodynamic approaches (Vienna RNAfold [Gruber et al., 2008], RNAstructure [Reuter and Mathews, 2010]) and a range of machine learning methods spanning multiple generations: from statistical learning approaches (CONTRAfold [Do et al., 2006], EternaFold [Wayment-Steele et al., 2022]) to deep learning architectures (UFold [Fu et al., 2022], MXFold2 [Sato et al., 2021]) and language models (RNA-FM [Chen et al., 2022]), allowing us to compare traditional energy-based algorithms against increasingly sophisticated data-driven approaches. Our **3° structure prediction** benchmarks encompass the latest advancements in 3D RNA modeling: AlphaFold3 [Abramson et al.], NuFold [Kagaya et al.], RhoFold+ [Shen et al.], RosettaFold2NA [Baek et al.], and trRosettaRNA [Wang et al.]. These models represent diverse methodological approaches, from end-to-end deep learning systems to hybrid methods that combine physics-based and machine learning approaches. Overall, this comprehensive selection of baselines allows us to systematically evaluate the strengths of different modeling paradigms across varying RNA types and prediction tasks. Additional details about baselines are provided in Appendix C.

### 3.4 Evaluation

For **fitness prediction**, the evaluation was primarily based on the Spearman’s rank correlation between the model predictions and experimental measurements, the Area Under the Curve (AUC) and the Matthews Correlation Coefficient (MCC). These metrics are complementary and were chosen to provide a comprehensive evaluation: Spearman correlation assesses the overall ranking of predictions, AUC measures the model’s ability to distinguish between functional and non-functional mutations, while MCC offers a balanced measure for potentially imbalanced datasets. To mitigate biases associated with uneven assay distributions across different RNA types, we calculated an average performance for each RNA type separately and then computed the overall performance as the mean of these RNA-type-level averages. This approach ensures that our results are robust and reflective of true model capabilities across varied biological categories. For the **2° structure prediction task**, we employed three standard metrics: F1-score, Area Under the Curve (AUC), and Mean Absolute Error (MAE). F1-score provides a balanced measure of precision and recall in identifying nucleotide pairings. AUC assesses the model’s ability to distinguish between paired and unpaired nucleotides. MAE offers a direct measure of prediction accuracy by quantifying the average magnitude of errors. For the **3° structure prediction task**, we employed TM score, ΔTM score, Watson-Crick intra-network fidelity (INF_WC_), and non-Watson-Crick INF (INF_NWC_), *ρ_TM_*, and *ρ_EC_*. TM-score captures overall fold correctness, while ΔTM-score captures predictive performance relative to the best scoring template structure. INF_WC_ and INF_NWC_ assess whether models correctly infer canonical and non-canonical base pairing. *ρ_TM_*, and *ρ_EC_* represent Spearman’s rank correlation with TM_train_and with the number of correct evolutionary couplings in the input alignment, respectively. In combination, these metrics gauge both global RNA fold and local base-pair fidelity, while ensuring equitable assessment despite distinct baseline training sets.

## 4 Results

### 4.1 Fitness prediction performance

#### Performance on all assays

The overall fitness prediction benchmark results (see Table 4) show Evo (2.0) and RNA-FM as the leading performers, with Spearman correlations of 0.276 and 0.217 respectively. These results indicate a leading capability for Evo 2.0 in predicting RNA fitness outcomes based on experimental data (statistical significance analysis is included in Appendix D.1). The relatively low scores across all models, particularly when compared to the stronger correlations reported for protein language models Notin et al. [2023], suggest substantial room for improvement and warrant deeper investigation. Several factors may contribute to this performance gap. A primary consideration is the limited availability of large-scale, diverse RNA datasets for model training compared to the abundance of protein sequence data. Additionally, there may be a potential misalignment between the training data used for these models and the taxonomical and functional distribution of our fitness landscapes. Lastly, differences in evolutionary conservation patterns between RNAs (especially non-coding RNAs) and proteins could also play a role, potentially affecting the models’ ability to capture fitness-relevant features.

**Table 4:** Fitness prediction overall performance. Average of Spearman’s rank correlation, AUC and MCC between model scores and experimental measurements on the full fitness prediction benchmark.

| Rank | Model name | Spearman | AUC | MCC |
| --- | --- | --- | --- | --- |
| 1 | Evo 2 | **0.276** | **0.636** | **0.212** |
| 2 | RNA-FM | 0.217 | 0.605 | 0.155 |
| 3 | RNAErnie | 0.189 | 0.598 | 0.154 |
| 4 | Evo 1.5 | 0.177 | 0.583 | 0.131 |
| 5 | Nucl. Transformer | 0.163 | 0.577 | 0.115 |
| 6 | RiNALMo | 0.155 | 0.582 | 0.122 |
| 7 | Evo 1 | 0.124 | 0.561 | 0.091 |
| 8 | GenSLM | 0.122 | 0.558 | 0.083 |

### Performance by RNA type

When examining performance by RNA type (Table A4), several models show specialized strengths across different RNA categories. RNA-FM achieves the highest correlations across all non coding RNAs including tRNA (0.464), Aptamer (0.190), and Ribozymes (0.201). On the other hand, the Evo family of models, and particularly Evo 2.0 have the best performances on mRNA-splicing (0.431) and mRNA coding (0.270) assays. These performance variations likely reflect the diverse training data of each model. RNA-FM’s particular strength with tRNAs, aptamers, and ribozymes aligns with its training on non-coding RNAs from RNAcentral, a database rich in these RNA types . The consistently strong performance of Evo models across different RNA types, and particularly in mRNAs, suggests their training approach may capture broader sequence-function relationships. These observations underscore the importance of targeted model training and selection based on the specific RNA type being studied. They also suggest that performance could potentially be improved by more tailored training data selection or by developing ensemble methods that leverage the strengths of different models for specific RNA types.

### Performance by mutation type

The fitness prediction results segmented by mutation type (Table A5) show that the Evo 2.0 model consistently outperform others across both single (0.230) and multiple (0.268) mutations. While these results establish Evo models as the current leaders in mutation effect prediction, the relatively low correlation values indicate substantial room for improvement in capturing RNA sequence-function relationships.

### Future Directions

To advance the field of RNA fitness prediction, several promising avenues of investigation emerge. First, developing models specifically trained on diverse RNA fitness landscapes could potentially improve performance by more closely aligning the training data with the prediction task. Additionally, incorporating RNA secondary structure predictions or experimental structure data into fitness prediction models may enhance their accuracy by capturing the important relationship between RNA structure and function. Comparing zero-shot performance with fine-tuned models could provide valuable insights into the generalizability of learned RNA features, potentially guiding future model development strategies. Lastly, exploring new architectural elements or pre-training objectives that better capture RNA-specific properties might lead to more robust and accurate predictions.

### 4.2 2° Structure prediction performance

#### Performance

The RNAGym structure prediction benchmark (Table 5) reveals interesting performance patterns across different RNA structure prediction methods for DMS and 2A3 chemical mapping data (see breakdown in Table A11). When considering overall performance, EternaFold demonstrates the strongest results in terms of F1-score and AUC getting 0.656 and 0.712 respectively. CONTRAfold follows closely with an F1-score of 0.653, then Vienna (0.644) and RNAstructure (0.643). This relatively tight performance distribution suggests that these more traditional RNA folding packages, while effective, may be approaching a methodological ceiling in their current framework. Among the more recent deep learning models, RNA-FM performs the best with respect to MAE (0.274). When comparing against RibonanzaNet to the other models, it underperforms for DMS chemical reactivity but does the best for 2A3 chemical reactivity (Table A12). 2A3 is a more unbiased signal for structure due to the reactivity being based on the 2’ hydroxyl instead of the nucleophilic nitrogens in adenines and cytosines. Since RibonanzaNet is trained with its own train-test split the results may be biased by data leakage.

**Table 5:** 2° structure prediction benchmark performance. F1-score, AUC and MAE between model predictions and experimental measurements (DMS and 2A3).

| Ranking | Model name | F1-score ↑ | AUC ↑ | MAE ↓ |
| --- | --- | --- | --- | --- |
| 1 | EternaFold | **0.656** | **0.712** | 0.338 |
| 2 | CONTRAFold | 0.653 | 0.708 | 0.353 |
| 3 | Vienna | 0.644 | 0.694 | 0.308 |
| 4 | RNAstructure | 0.643 | 0.692 | 0.312 |
| 5 | RNA-FM | 0.601 | 0.645 | **0.274** |
| 6 | UFold | 0.574 | 0.611 | 0.324 |
| 7 | MXfold2 | 0.518 | 0.529 | 0.446 |

### Future Directions

Our evaluation reveals that current zero-shot RNA structure prediction methods perform remarkably similarly, with only small differences separating their effectiveness. To overcome this limit, significant advances will likely come from supervised learning approaches that can leverage experimental data more effectively. To facilitate this progress, we provide a carefully curated nonredundant training dataset that researchers can use to develop and benchmark supervised methods without concerns of sequence redundancy between training and test sets. By establishing this resource, we aim to enable fair evaluation of model generalization capabilities and accelerate the development of next-generation RNA structure prediction methods that can significantly outperform current approaches while maintaining robustness across diverse RNA families.

### 4.3 3° Structure prediction performance

#### Performance

The overall tertiary structure benchmark results (see Table 6) show that NuFold leads for monomeric RNAs, with a top TM score of 0.393, while AlphaFold3 emerges as the best performer on complexes (TM = 0.381). NuFold’s correlations with training-set similarity (*ρ_TM_* = 0.74) and evolutionary couplings (*ρ_EC_* = 0.67) are also the highest for monomers, implying that it leverages both memorized structural priors and covariation signals effectively. Notably, the top monomeric models exhibit modest ΔTM values (Δ*TM ≈ −*0.15), suggesting that, despite relying on different training sets they learn comparable structural models with room for improvement when generalizing beyond memorized templates. AlphaFold3 tops total Watson–Crick (INF_WC_=0.83) and non-Watson-Crick interactions (INF_NWC_=0.26), followed closely by NuFold, RoseTTAFold2NA, and trRosettaRNA. The notably lower scores for non-Watson-Crick interactions underscores a significant gap in RNA structure prediction, highlighting an area where future advances are urgently needed. For RNA complexes, AlphaFold3 outperforms RoseTTAFold2NA, achieving a TM score of 0.380 vs. 0.167. Interestingly, AlphaFold3’s performance on complexes matches or exceeds its performance on monomers despite less homology to training (Δ*TM* = −0.13), highlighting its robust handling of multi-chain interfaces. Moreover, it shows higher correlation with evolutionary couplings (*ρ_EC_* = 0.77) than with training-set similarity (*ρ_TM_* = 0.55), suggesting that it can leverage covariation signals effectively even in these more challenging contexts. The pronounced drop-off in performances for RoseTTAFold2NA indicates that complexes remain difficult for certain architectures.

**Table 6:** RNAGym - 3° structure prediction results. NuFold leads monomers and AlphaFold3 leads complexes with TM scores of about 0.4, reflecting moderate prediction accuracy despite structural diversity. Monomers shown at top, complexes at bottom.

| Rank | Model name | TM | ΔTM | $\rho_{TM}$ | $\rho_{EC}$ | INF<sub>WC</sub> | INF<sub>NWC</sub> |
| --- | --- | --- | --- | --- | --- | --- | --- |
| 1 | NuFold | **0.393** | **-0.151** | **0.74** | **0.67** | 0.78 | 0.19 |
| 2 | AlphaFold3 | 0.386 | -0.167 | 0.71 | 0.60 | **0.83** | **0.26** |
| 3 | trRosettaRNA | 0.382 | -0.152 | 0.60 | 0.58 | 0.75 | 0.12 |
| 4 | RhoFold+ | 0.367 | -0.172 | 0.72 | 0.62 | 0.50 | 0.09 |
| 5 | RoseTTAFold2NA | 0.365 | -0.154 | 0.67 | 0.55 | 0.74 | 0.21 |
| 1 | AlphaFold3 | **0.381** | **-0.130** | **0.55** | **0.77** | **0.86** | **0.37** |
| 2 | RoseTTAFold2NA | 0.168 | -0.294 | 0.24 | 0.55 | 0.18 | 0.00 |

### Future directions

These results indicate promising yet still limited zero-shot accuracy for RNA tertiary structure prediction, Although TM and ΔTM scores reveal room for improvement, different models demonstrate unique advantages: NuFold excels at template modeling and evolutionary coupling extraction (ΔTM, *ρ_TM_*, *ρ_EC_*), while AlphaFold3 leads on local basepair network fidelity (INF_WC_, INF_NWC_) and complexes. The poor performance of all models on non-Watson-Crick interactions highlights a pressing need for new approaches. Harnessing these complementary strengths in a unified model will be instrumental to achieving highly accurate 3D RNA predictions. The centralized RNAGym benchmark accelerates progress towards this goal by enabling the community to more transparently compare the strengths and weaknesses of new methods as they emerge.

## 5 Resources

### Codebase

We open source (MIT License) all resources curated for the RNAGym benchmark via our GitHub repository. In particular, we consolidate of the numerous RNA structure and fitness prediction models discussed in Appendix C and make them available via a common interface. This resource aims to provide researchers with robust tools, reducing the technical barrier to entry for conducting advanced RNA analysis and facilitating the reproducibility of results.

### Processed datasets

We have made available all datasets used in our fitness and structure prediction benchmarks, including both raw and processed versions, as detailed in Section 3.2 at huggingface, and on our GitHub repository. To enhance the utility of our benchmarks, we have included several additional components. For the fitness benchmark, where available, we provide tertiary structure PDB files and multiple sequence alignments for the relevant protein families. For the 2° structure benchmark, we have clustered the sequences at a 40% sequence identity cutoff and provide both the representative clusters as well as the clustering results from mmseq2. From the clustering we created a clean non-redundant 20-80 train-test datasplit. We have mapped all sequences in the train and test set to similar RNA sequences found in the RCSB, PseudoBase [van Batenburg et al., 2000], and bpRNA [Danaee et al., 2018] databases, providing easy access to the rich annotations contained in these databases. Furthermore, to support researchers interested in supervised learning approaches, we offer training datasets for both the fitness and secondary structure prediction tasks (Appendix B).

## 6 Conclusion

RNAGym addresses the significant gap in large-scale benchmarks for the robust evaluation of models tailored for RNA structure prediction and fitness assessment. It enables the direct comparison of methods across several dimensions of interest (e.g., RNA type, mutation type). We anticipate that the RNAGym benchmarks and the accompanying data assets we release to the public will serve as invaluable resources for the Machine Learning and Computational Biology communities. We plan to continually update the benchmarks as new data and baseline models become available.

## Acknowledgments

R.W. is supported by the EPSRC Centre for Doctoral Training in Health Data Science (EP/S02428X/1). D.S.M. and P.N. are supported by a Chan Zuckerberg Initiative Award (Neurodegeneration Challenge Network, CZI2018-191853). This research was conducted in part on the O2 High Performance Compute Cluster, supported by the Research Computing Group, at Harvard Medical School. The authors also acknowledge the support of Dr. Scott T. Aoki and the Indiana University Pervasive Technology Institute for providing supercomputing and storage resources that have contributed to the research results reported within this paper.

## Appendix

### A Dataset collection

#### Prioritization of studies for expert review

Coding mRNA datasets were sourced from ProteinGym when nucleotide-level deep mutational scanning information was available [Notin et al., 2023].

The selection process for prioritizing noncoding RNA studies for expert review was structured as follows. We initiated with a targeted PubMed search, utilizing specific queries to ensure a comprehensive capture of relevant literature:

1. **Deep Mutational Scan:** (deep[All Fields] OR comprehensive[All Fields]) AND (mutational[All Fields] OR muta*) AND (scan OR scans OR scanning) AND RNA
2. **Saturation Mutagenesis:** (saturat* muta*) AND RNA
3. **MAVE:** (“Multiplex* assay” AND “variant*”) AND RNA
4. **MPRA:** (“MPRA” OR “Massively parallel reporter assay*”) AND RNA NOT protein
5. **Other:** (“Fitness Landscape” AND muta*) AND RNA

These initial searches proved to be either overly restrictive or too broad, which complicated the manual screening process. Ultimately, this approach resulted in the identification of only 20 primary articles. To enrich our review, an additional 10 articles were identified by scraping references from other pertinent studies, including those cited in previous research such as [Sumi et al., 2024, Nguyen et al., 2024], and RNA-related datasets from studies like ProteinGym [Notin et al., 2023].

Hypothesizing that our initial search strategy may have missed relevant studies, we conducted a comprehensive PubMed search using the following query:

#### Broad Search

(deep OR comprehensive OR MPRA OR multiplex assay OR massively parallel OR landscape OR saturation) AND (muta* OR variant OR variants) AND (scan OR scans OR screen OR landscape OR assay) AND (RNA OR ribozyme* OR microRNA OR miRNA OR siRNA OR snoRNA OR tRNA OR lncRNA OR (RNA AND aptamer) OR circRNA)

#### **A.1** Literature pre-screening with LLM

The prior search yielded an overwhelming 11,635 results. To efficiently handle this volume, we utilized a large language model (LLM), specifically GPT4-0125-preview, for secondary screening. We adapted a recent prompting approach designed for systematic review screening [Cao et al., 2024]. The LLM was instructed with clear study objectives and specific inclusion/exclusion criteria, effectively narrowing down the pool to fewer than 500 articles, thereby making manual curation manageable. To enhance the sensitivity of this process, the LLM’s prompt was refined using an initial set of 30 positively identified articles as a control group. This novel use of LLMs for data extraction markedly improved our capacity to pinpoint relevant studies. Consequently, we were able to incorporate an additional 22 studies into our initial screen, resulting in a total of 52 studies ready for manual expert review.

We used the following prompt to pre-screen relevant studies during our extensive literature search:

"The goal of this study is to create a benchmark that contains RNA deep mutational scanning or fitness landscape datasets. We are generating these datasets to benchmark RNA fitness prediction algorithms, and need our datasets we evaluate to have information on RNA mutants/variants and their relative ’fitness’.

The following is an excerpt of two sets of criteria. A study is considered included if it meets all the inclusion criteria. If a study meets any of the exclusion criteria, it should be excluded. Here are the two sets of criteria:

Inclusion Criteria (all must be fulfilled): 1. Studies involve RNA. We are also interested in RNA subclasses such as Ribozyme, lncRNA, tRNA, rRNA, microRNA (miRNA), Aptamer, Riboswitch, mRNA 2. Studies report on fitness prediction. Other terms for fitness prediction can include deep mutational scans, comprehensive multiplex assays, or comprehensive fitness landscapes, among others 3. Studies with greater than 100 experimental measurements 4. Studies that report on mutant fitness through reporter assays, bulk RNA-sequencing, single-cell RNA sequencing assay, fluorescence in-situ hybridization (FISH) assay, flow cytometry assay, imaging mass cytometry assay, evolution of ligands by exponential enrichment assay, single cell imaging, multiplexed fluorescent antibody imaging, binding assays, cell proliferation assay, splicing assays, survival assessment assay selection types, or similar. 5. Studies that report on enzymatic activity, binding affinity, stability, fluorescence, proliferation selection assays, or similar assays. 6. The study must be primary research and generate a novel dataset

Exclusion Criteria (if any met then exclude): 1. Studies only reporting on protein mutational scans, with no relevance or mention of RNA being mutated 2. Studies that do not focus on fitness quantification 3. Review articles (systematic reviews, case reports, case series, etc.) or other non-primary research sources.

##### Instructions

We now assess whether the paper should be included in the systematic review by evaluating it against each and every predefined inclusion and exclusion criterion. First, we will reflect on how we will decide whether a paper should be included or excluded. Then, we will think step by step for each criteria, giving reasons for why they are met or not met. Studies that may not fully align with the primary focus of our inclusion criteria but provide data or insights potentially relevant to our review deserve thoughtful consideration. Given the nature of abstracts as concise summaries of comprehensive research, some degree of interpretation is necessary. Our aim should be to inclusively screen abstracts, ensuring broad coverage of pertinent studies while filtering out those that are clearly irrelevant. We will conclude by outputting (on the very last line) ’XXX’ if the paper warrants exclusion, or ’YYY’ if inclusion is advised or uncertainty persists. We must output either ’XXX’ or ’YYY’.

Title and Abstract in investigation:

Title: #Insert title of study#

Abstract: #Insert abstract of paper#"

#### Expert review process

The process for accepting a paper involved several steps to ensure the quality and relevance of the data. First, we checked whether the data was openly available and could be integrated into our benchmark. If data was not accessible, study authors were contacted.

Next, we used the following inclusion and exclusion criteria during our through expert review process:

#### Inclusion Criteria

- Assay must focus on RNA
- Assay must have at least 100 experimental variants tested, with a sufficiently wide dynamic range
- Assay must be relevant to fitness prediction, and report on mutant fitness
- Assay must only focus on substitutions, not insertions or deletions

#### Exclusion Criteria

- Assays focusing on DNA or Proteins
- Assays that are not primary research
- Assays with mutants of varying lengths

### B Datasets details

#### B.1 Fitness assays

##### References

An exhaustive list of the publications from which the assays included our fitness benchmark originated from is provided in Table B.1 and Table B.1. Our final processed datasets all have a consistent format with the same 3 fields across: "Mutant" (mutation triplets), "Sequence" (mutated sequence), and "DMS score" (experimental measurement). We also corrected the *directionality* of each measured experimental phenotype, to ensure that higher DMS scores always translate to higher fitness across assays.

##### Licenses

All fitness assays were licensed under CC-BY 4.0 (https://creativecommons.org/licenses/by/4.0/), or the ACS AuthorChoice Usage Agreement (https://pubs.acs.org/page/policy/authorchoice_termsofuse.html).

##### Cross-validation splits

For users interested in supervised RNA fitness prediction, we provide two types of cross-validation splits:

• **Random**: a random 80%-20% train-test split;

• **Minimum similarity**: a 80%-20% split in which we minimized the sequence similarity between training and validation RNA sequences.

#### B.2 Structure prediction dataset

We constructed the data for our structure prediction challenge using the data from the Ribonanza Challenge hosted on Kaggle (https://www.kaggle.com/competitions/stanford-ribonanza-rna-folding/data) and RMDB [Cordero et al., 2012]. The data contains RNA chemical reacitivty at various conditions (Table 2). For ease of use we created a dataset of only in vitro chemical mapping data involving 2A3 and DMS. While we do not include the data such as those collected with ligands in our clean train-test data, it is available. To ensure the integrity of our dataset and prevent data leakage, we clustered the sequences with 40% sequence identity and created a 20-80 train-test split. The final dataset consists of 901*k* reactivity profiles for 583*k* unique sequences and 100*M* nucleotides. This extensive dataset underpins the robustness and comprehensive nature of our RNA structure prediction challenge. The original data from the challenge is made available under a CC-BY 4.0 license.

### C Baselines

#### C.1 RNA fitness prediction models

Our fitness prediction benchmarks currently include the following 5 baselines:

• **RiNALMo** [Penić, et al., 2024] is a 650 million parameter RNA language model trained on 36 million non-coding RNA sequences, achieving high performance in RNA structural and functional prediction tasks. Variants were scored using the masked marginal scoring strategy from the ESM sequence modeling framework.

• **Evo** [Nguyen, et al., 2024] is a 7 billion parameter model trained on 2.7M prokaryotic and phage genomes to generate DNA sequences using a context length of 8k (further extended to131k tokens). It is based on StripedHyena, a deep signal processing architecture designed to improve efficiency and quality over the prevailing Transformer architecture. We use the 8k version of the model, refered as Evo 1. The recently released Evo 1.5 builds upon Evo 1 by increasing the pretraining data from 300 billion to 450 billion tokens Merchant et al. [2024]. We also include the EVO2 (7B) model Brixi et al. [2025], EVO2 is trained on over 9.3T tokens at a single nucleotide resolution.

• **RNA-FM** [Chen, et al., 2022] is a RNA foundation model based on the BERT language model architecture. It was trained on 23 million unlabeled non-coding RNA sequences from over 800,000 species, collected from RNA-central. Variants were scored using the wild-type marginal scoring strategy from the ESM sequence modeling framework, which RNA-FM builds upon.

• **GenSLM** [Zvyagin, et al., 2023] includes an autoregressive RNA language model with codon-level tokenization, along with stable diffusion, to model genome-scale interactions and predict SARS-CoV-2 evolution. It was trained on 110 million prokaryotic coding sequences from BV-BRC. Variant sequence likelihoods were scored using the 2.5 billion parameter language model.

• **RNAErnie** [Wang, et al., 2024b] is a 12-layer transformer-based RNA language model pre-trained on 23 million ncRNA sequences using masked language modeling. With 105 million trainable parameters, it achieves high performance in RNA sequence classification, RNA–RNA interaction, and RNA secondary structure prediction.

• **Nucleotide Transformer** [Dalla-Torre et al., 2023] is a 2.5B parameter model trained on 850 species. The 2.5b-multi-species version was used.

#### C.2 RNA 2° structure prediction models

Our 2° structure prediction benchmarks currently include the following 7 baselines:

• **EternaFold** [Wayment-Steele, et al., 2020] is built on the principles derived from the Eterna massive open online game, where players design RNA sequences that fold into target shapes. This model incorporates crowd-sourced insights from thousands of players to refine its algorithms, significantly enhancing its ability to predict RNA structures under varied environmental conditions and complexities.

• **CONTRAfold** [Do et al., 2006] is a machine learning-based RNA secondary structure prediction model that utilizes conditional log-linear models for structure inference.

• **Vienna** [Gruber et al., 2008] is one of the most widely used RNA secondary structure prediction tools. It employs dynamic programming algorithms based on thermodynamic models to accurately predict RNA secondary structures, including handling pseudoknotted structures as extensions.

• **RNAstructure** [Reuter and Mathews, 2010] is a model designed for the prediction and analysis of RNA secondary structures. It is known for its dual ability to use either thermodynamic or machine learning-based methods to predict RNA folding patterns.

• **RNA-FM** [Chen et al., 2022] as described above was also used for RNA structure prediction by utilizing its downstream second-order structure prediction module.

• **UFold** [Fu et al., 2022] is a deep learning approach that employs a U-Net architecture to predict RNA secondary structures.

• **Mxfold2** [Sato et al., 2021] integrates deep neural networks with thermodynamic parameters for RNA structure prediction. Using a max-margin framework with thermodynamic regularization, it combines folding scores from neural networks with Turner’s energy parameters.

#### C.3 RNA 3° structure prediction models

Our tertiary structure prediction benchmarks currently include the following 5 baselines:

• **AlphaFold3** [Abramson et al.] extends the AlphaFold framework to RNA, predicting full 3D coordinates using a diffusion-based architecture trained on protein and nucleic acid data. It captures interchain interactions, supporting both RNA monomers and complexes.

• **NuFold** [Kagaya et al.] is an end-to-end deep learning model dedicated to RNA, adapting iterative refinement techniques from protein structure prediction. It incorporates base-centered representations to accurately capture local RNA geometry.

• **RhoFold+** [Shen et al.] offers an automated, end-to-end platform for RNA 3D structure prediction that integrates the RNA-FM language model and iterative geometry-aware refinement.

• **RoseTTAFold2NA** [Baek et al.] extends the RoseTTAFold framework to handle RNA and protein–RNA complexes. By jointly refining sequence features, residue-pair distances, and cartesian coordinates, it models RNA folds and interfaces in a single pipeline.

• **trRosettaRNA** [Wang et al.] leverages a transformer-based network to predict intra-nucleotide distances and torsion angles, followed by refinement using Rosetta’s energy minimization. This hybrid approach combines deep-learning potentials and physics-based energy terms to generate accurate RNA structures.

### D Detailed experimental results

#### D.1 Detailed fitness prediction performance

We report the assay-level Spearman performance, across all assays in the RNAGym fitness prediction benchmark in Fig D.1, aggregated performance by RNA type in Table A4, and by mutational depth in Table A5. Additionally, we report Taxa level Spearman performance in Table A6, MCC performance in Table A8, and AUC performance in Table A7 in for our coding fitness prediction benchmark.

We also report the statistical significance for the relative Spearman performance by RNA type in Table A9. We follow the same methodology as in ProteinGym [Notin et al., 2023] and assess statistical significance by computing the non-parametric bootstrap standard error of the difference between the Spearman performance of a given model and that of the best overall model.

#### D.2 Secondary structure prediction

##### Chemical mapping type

We provide a performance breakdown for the different models against our chemical mapping test set in Table A11. In order to create a fair comparison for RibonanzaNet, we created a separate test set consisting of the intersection of RibonanzaNet test set [He et al., 2024] with our test split. We measured the performance of all the models against this smaller test set in Table A12.

##### Pseudoknots

We mapped RNA sequences in our evaluation set to pseudoknot annotations from PseudoBase [van Batenburg et al., 2000] (358 test sequences mapped), and report the corresponding global F1 score and crossed pair F1 score (Table A13). Out of our various structure prediction baselines, only RNAstructure is able to score pseudoknots.

### E Limitations

Experimentally assaying RNA fitness, while resource-intensive, provides critical insights that help advance our understanding of RNA function. However, such experimental assays may not always accurately mimic the cellular environment, which can lead to variations between observed in vitro results and actual in vivo functionality. Chemical mapping experiments using dimethyl sulfate (DMS) offer valuable data on RNA secondary structures by identifying accessible adenine and cytosine bases that interact with DMS. This technique, although powerful for revealing the in-vivo-like structure of RNA in a relatively high-throughput manner, has its limitations. DMS mapping can be affected by incomplete coverage, as it primarily marks only two of the four nucleotide types. Additionally, the resolution of DMS mapping might not always distinguish closely spaced structural features, potentially obscuring important details about RNA folding and interaction sites. The accuracy of predictions from DMS data also heavily depends on the computational tools used to interpret the chemical reactivity patterns, necessitating ongoing improvements in both experimental and analytical methodologies to enhance the precision and utility of RNA structural studies. There is also sampling bias for the RNA families that were assayed using high-throughput fitness assays, with not all families equally represented. The majority of fitness assays were focused on ribozymes while other RNA families such as mRNA and tRNA had far fewer available datasets to evaluate.

**Figure 2:**
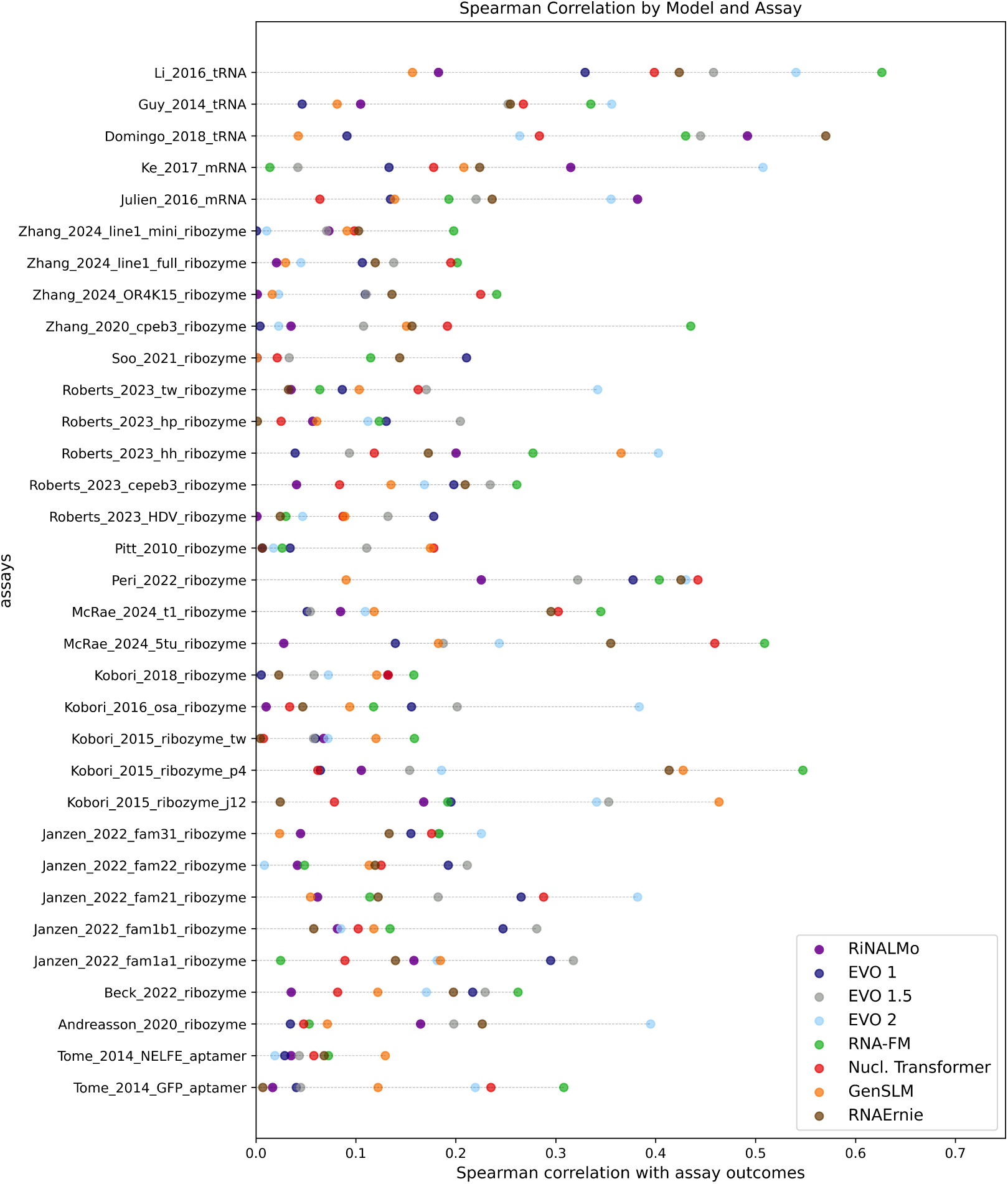
RNAGym fitness prediction benchmark - Detailed performance for non-coding and splicing assays. Spearman’s rank correlation between model predictions and experimental values for non coding and splicing assays in the RNAGym fitness prediction benchmark.

**Figure 3:**
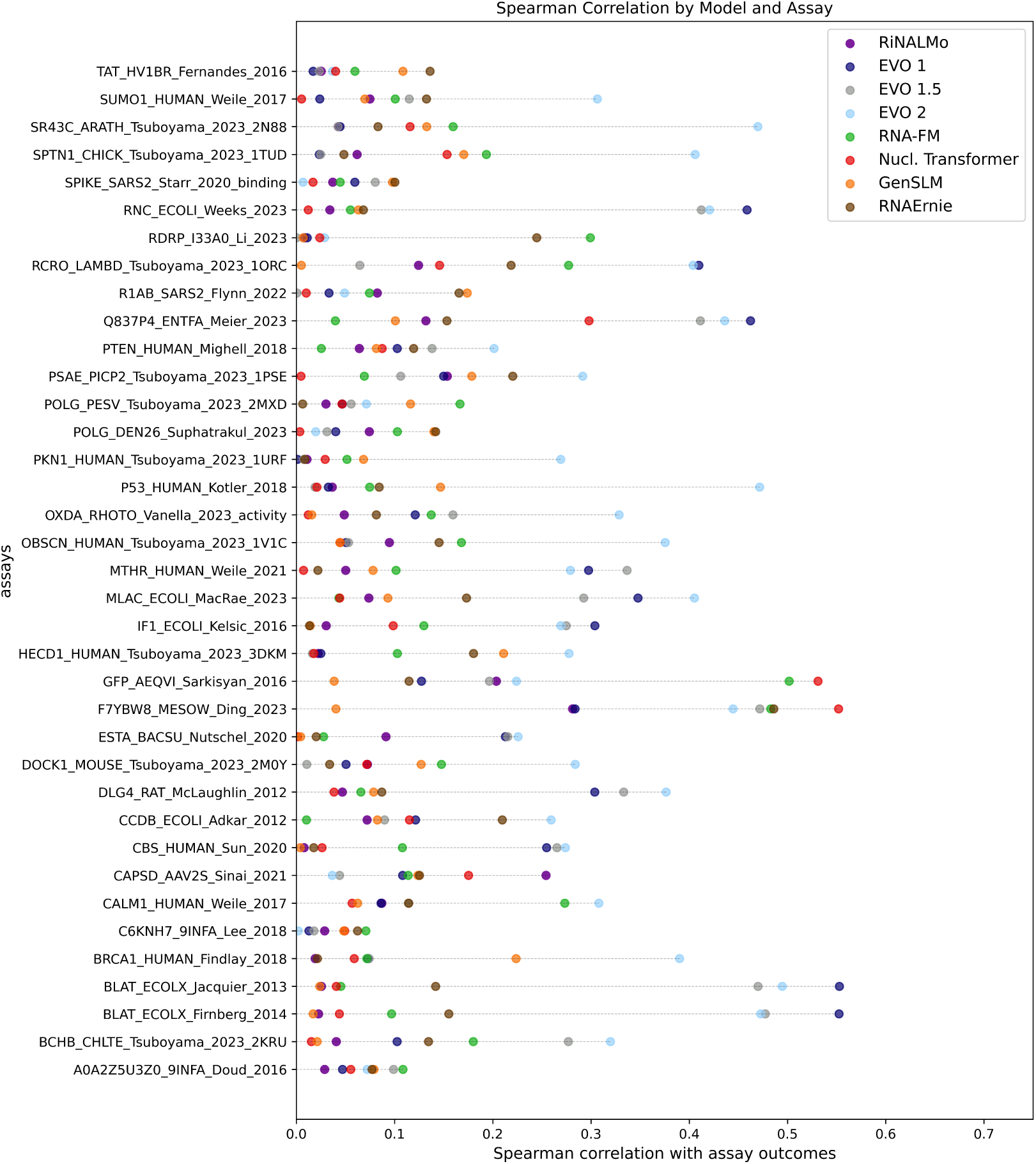
RNAGym fitness prediction benchmark - Detailed performance for coding assays. Spearman’s rank correlation between model predictions and experimental values for coding assays in the RNAGym fitness prediction benchmark.

### F Societal impact

The advancement of RNA models holds transformative potential across a spectrum of applications. By accurately predicting RNA structure and fitness, researchers can unlock new therapies by targeting previously intractable genetic conditions, enhance crop resilience through agricultural biotechnology, and even engineer microbial systems for cleaner energy production. The creation of benchmarks like RNAGym is crucial in this endeavor, as they drive the field forward by setting standards for model performance and fostering innovation through competition and collaboration. However, as it is the case for any approach that facilitates the development of novel biological sequences for good, the potential misuse of these technologies to create harmful biological agents cannot be ignored [Urbina et al., 2022]. It is imperative to proceed with a careful framework that promotes secure use, ethical guidelines, and synthesis monitoring [Baker and Church, 2024] to mitigate risks associated with dual-use capabilities. Ultimately, benchmarks like RNAGym not only validate the effectiveness of emerging RNA models but also, by highlighting the methods leading to step-change performance improvements, encourage their integration into real-world applications, ensuring that these innovations contribute positively to society.

### G Compute resources

In our benchmarking work for RNA fitness prediction and structure analysis, we primarily rely on GPUs as hardware accelerators to handle the computationally intensive tasks involved. We estimate our compute budget for both fitness and structure prediction benchmarks to approximately 60 V100 GPU days.

**Table A1:** RNAGym fitness prediction data. We developed our noncoding and splicing fitness prediction benchmark by curating and processing 33 assays from 19 publications.

| Title | Year | RNA type | # Assays | Reference | License |
| --- | --- | --- | --- | --- | --- |
| Saturation mutagenesis reveals manifold determinants of exon definition | 2017 | mRNA - splicing | 1 | [Ke et al., 2017] | CC BY 4.0 |
| Fitness landscape of a dynamic RNA structure | 2021 | ribozyme | 1 | [Soo et al., 2021] | CC BY 4.0 |
| Comprehensive sequence-to-function mapping of cofactor-dependent RNA catalysis in the glmS ribozyme | 2020 | ribozyme | 1 | [Andreasson et al., 2020] | CC BY 4.0 |
| High-throughput assay and engineering of self-cleaving ribozymes | 2015 | ribozyme | 3 | [Kobori et al., 2015] | CC BY 4.0 |
| Identification of the determinants of tRNA function and susceptibility to rapid tRNA decay by high-throughput in vivo analysis | 2014 | tRNA | 1 | [Guy et al., 2014] | CC BY 4.0 |
| Deep sequencing analysis of aptazyme variants based on a pistol ribozyme | 2018 | ribozyme | 1 | [Kobori et al., 2017] | ACS Author-Choice Usage Agreement |
| Accurate inference of the full base-pairing structure of RNA by deep mutational scanning and covariation-induced deviation of activity. | 2020 | ribozyme | 1 | [Zhang et al., 2019] | CC BY 4.0 |
| Rapid Construction of Empirical RNA Fitness Landscapes | 2010 | ribozyme | 1 | [Pitt and Ferré-D'Amaré, 2010] | CC-BY 4.0 |
| Dynamic RNA Fitness Landscapes of a Group I Ribozyme during Changes to the Experimental Environment | 2022 | ribozyme | 1 | [Peri et al., 2022] | CC-BY 4.0 |
| RNA sequence to structure analysis from comprehensive pairwise mutagenesis of multiple self-cleaving ribozymes | 2023 | ribozyme | 5 | [Roberts et al., 2023] | CC-BY 4.0 |
| Pairwise and higher-order genetic interactions during the evolution of a tRNA | 2018 | tRNA | 1 | [Domingo et al., 2018] | CC-BY 4.0 |
| Emergent properties as by-products of prebiotic evolution of aminoacylation ribozymes | 2022 | ribozyme | 5 | [Janzen et al., 2022] | CC-BY 4.0 |
| Predicting higher-order mutational effects in an RNA enzyme by machine learning of high-throughput experimental data | 2022 | ribozyme | 1 | [Beck et al., 2022] | CC-BY 4.0 |
| The fitness landscape of a tRNA gene | 2016 | tRNA | 1 | [Li et al., 2016] | CC-BY 4.0 |
| Comprehensive analysis of RNA-protein interactions by high-throughput sequencing-RNA affinity profiling | 2014 | aptamer | 2 | [Tome et al., 2014] | CC-BY 4.0 |
| The complete local genotype-phenotype landscape for the alternative splicing of a human exon | 2016 | mRNA - splicing | 1 | [Julien et al., 2016] | CC-BY 4.0 |
| Minimal twister sister-like self-cleaving ribozymes in the human genome revealed by deep mutational scanning. | 2024 | ribozyme | 3 | [Zhang et al., 2024] | CC-BY 4.0 |
| Cryo-EM structure and functional landscape of an RNA polymerase ribozyme | 2024 | ribozyme | 2 | [McRae et al., 2024] | CC-BY 4.0 |
| High-Throughput Mutational Analysis of a Twister Ribozyme | 2016 | ribozyme | 1 | [Kobori and Yokobayashi, 2016] | CC-BY 4.0 |

**Table A2:** RNAGym coding fitness prediction data. We developed our coding fitness prediction benchmark by curating and processing 37 assays from 27 publications.

|  |  |  |  |  |  |
| --- | --- | --- | --- | --- | --- |
| A Comprehensive, High-Resolution Map of a Gene’s Fitness Landscape | 2014 | mRNA | 1 | [Firnberg et al., 2014] | CC-BY 4.0 |
| Capturing the mutational landscape of the beta-lactamase TEM-1 | 2013 | mRNA | 1 | [Jacquier et al., 2013] | CC-BY 4.0 |
| Protein model discrimination using mutational sensitivity derived from deep sequencing | 2012 | mRNA | 1 | [Adkar et al., 2012] | CC-BY 4.0 |
| RNA structural determinants of optimal codons revealed by MAGE-Seq | 2016 | mRNA | 1 | [Kelsic et al., 2016] | CC-BY 4.0 |
| Fitness and functional landscapes of the E. coli RNase III gene rnc | 2023 | mRNA | 1 | [Weeks and Ostermeier, 2023] | CC-BY 4.0 |
| Protein design using structure-based residue preferences | 2023 | mRNA | 1 | [Ding et al., 2024] | CC-BY 4.0 |
| Deep mutational scan of a drug efflux pump reveals its structure–function landscape | 2023 | mRNA | 1 | [Meier et al., 2023] | CC-BY 4.0 |
| Protein-protein interactions in the Mla lipid transport system probed by computational structure prediction and deep mutational scanning | 2023 | mRNA | 1 | [MacRae et al., 2023] | CC-BY 4.0 |
| Systematically Scrutinizing the Impact of Substitution Sites on Thermostability and Detergent Tolerance for Bacillus subtilis Lipase A | 2020 | mRNA | 1 | [Nutschel et al., 2020] | CC-BY 4.0 |
| Local fitness landscape of the green fluorescent protein | 2016 | mRNA | 1 | [Sarkisyan et al., 2016] | CC-BY 4.0 |
| The spatial architecture of protein function and adaptation | 2012 | mRNA | 1 | [McLaughlin Jr et al., 2012] | CC-BY 4.0 |
| Understanding Activity-Stability Tradeoffs in Biocatalysts by Enzyme Proximity Sequencing | 2023 | mRNA | 1 | [Vanella et al., 2024] | CC-BY 4.0 |
| A Systematic p53 Mutation Library Links Differential Functional Impact to Cancer Mutation Pattern and Evolutionary Conservation | 2018 | mRNA | 1 | [Kotler et al., 2018] | CC-BY 4.0 |
| Accurate classification of BRCA1 variants with saturation genome editing | 2018 | mRNA | 1 | [Findlay et al., 2018] | CC-BY 4.0 |
| Deep Mutational Scanning of SARS-CoV-2 Receptor Binding Domain Reveals Constraints on Folding and ACE2 Binding | 2020 | mRNA | 1 | [Starr et al., 2020] | CC-BY 4.0 |
| Deep mutational scanning of hemagglutinin helps predict evolutionary fates of human H3N2 influenza variants | 2018 | mRNA | 1 | [Lee et al., 2018] | CC-BY 4.0 |
| Mega-scale experimental analysis of protein folding stability in biology and design | 2023 | mRNA | 10 | [Tsuboyama et al., 2023] | CC-BY 4.0 |
| A framework for exhaustively mapping functional missense variants | 2017 | mRNA | 2 | [Weile et al., 2017] | CC-BY 4.0 |
| A Saturation Mutagenesis Approach to Understanding PTEN Lipid Phosphatase Activity and Genotype-Phenotype Relationships | 2018 | mRNA | 1 | [Mighell et al., 2018] | CC-BY 4.0 |
| A proactive genotype-to-patient-phenotype map for cystathionine beta-synthase | 2020 | mRNA | 1 | [Sun et al., 2020] | CC-BY 4.0 |

**Table A3:** RNAGym coding fitness prediction data (continued). We developed our coding fitness prediction benchmark by curating and processing 37 assays from 27 publications.

|  |  |  |  |  |  |
| --- | --- | --- | --- | --- | --- |
| Shifting landscapes of human MTHFR missense-variant effects | 2021 | mRNA | 1 | [Weile et al., 2021] | CC-BY 4.0 |
| Functional analysis of flavivirus replicase by deep mutational scanning of dengue NS5 | 2023 | mRNA | 1 | [Suphatrakul et al., 2023] | CC-BY 4.0 |
| Deep mutational scanning reveals the functional constraints and evolutionary potential of the influenza A virus PB1 protein | 2023 | mRNA | 1 | [Li et al., 2023] | CC-BY 4.0 |
| Comprehensive fitness landscape of SARS-CoV-2 Mpro reveals insights into viral resistance mechanisms | 2022 | mRNA | 1 | [Flynn et al., 2022] | CC-BY 4.0 |
| Accurate Measurement of the Effects of All Amino-Acid Mutations on Influenza Hemagglutinin | 2016 | mRNA | 1 | [Doud and Bloom, 2016] | CC-BY 4.0 |
| Functional Segregation of Overlapping Genes in HIV | 2016 | mRNA | 1 | [Fernandes et al., 2016] | CC-BY 4.0 |
| Generative AAV capsid diversification by latent interpolation | 2021 | mRNA | 1 | [Sinai et al., 2021] | CC-BY 4.0 |

**Table A4:**
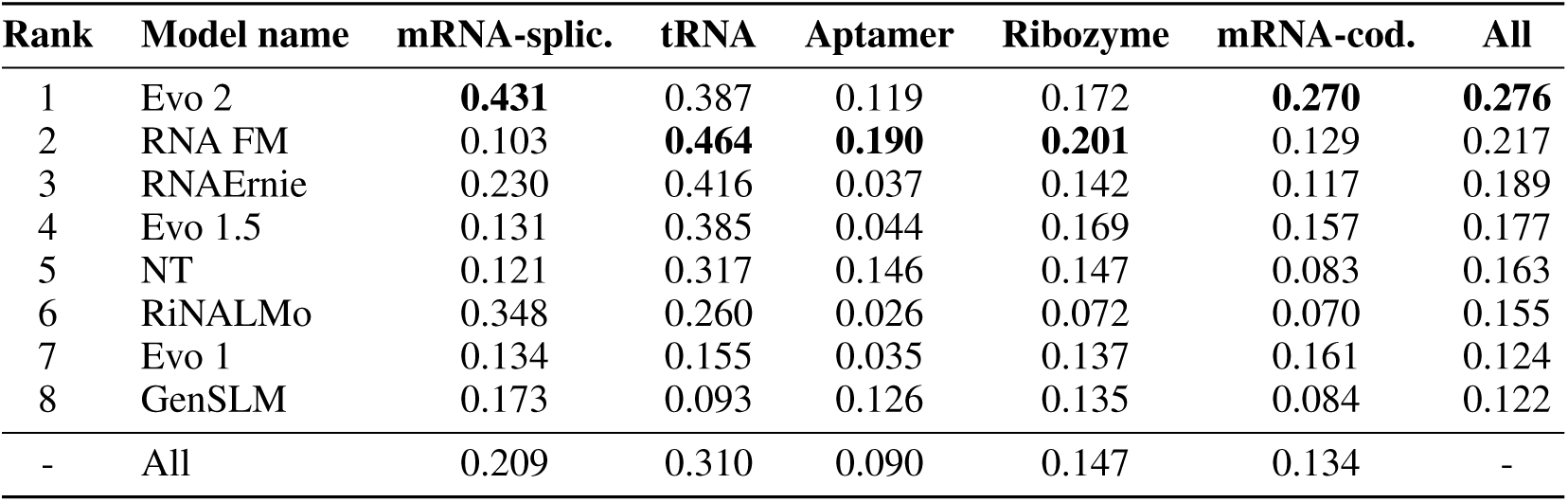
RNAGym - Fitness prediction by RNA type. Average of Spearman’s rank correlation between model scores and experimental measurements by RNA type and overall.

| Rank | Model name | mRNA-splic. | tRNA | Aptamer | Ribozyme | mRNA-cod. | All |
| --- | --- | --- | --- | --- | --- | --- | --- |
| 1 | Evo 2 | **0.431** | 0.387 | 0.119 | 0.172 | **0.270** | **0.276** |
| 2 | RNA FM | 0.103 | **0.464** | **0.190** | **0.201** | 0.129 | 0.217 |
| 3 | RNAErnie | 0.230 | 0.416 | 0.037 | 0.142 | 0.117 | 0.189 |
| 4 | Evo 1.5 | 0.131 | 0.385 | 0.044 | 0.169 | 0.157 | 0.177 |
| 5 | NT | 0.121 | 0.317 | 0.146 | 0.147 | 0.083 | 0.163 |
| 6 | RiNALMo | 0.348 | 0.260 | 0.026 | 0.072 | 0.070 | 0.155 |
| 7 | Evo 1 | 0.134 | 0.155 | 0.035 | 0.137 | 0.161 | 0.124 |
| 8 | GenSLM | 0.173 | 0.093 | 0.126 | 0.135 | 0.084 | 0.122 |
| - | All | 0.209 | 0.310 | 0.090 | 0.147 | 0.134 | - |

**Table A5:**
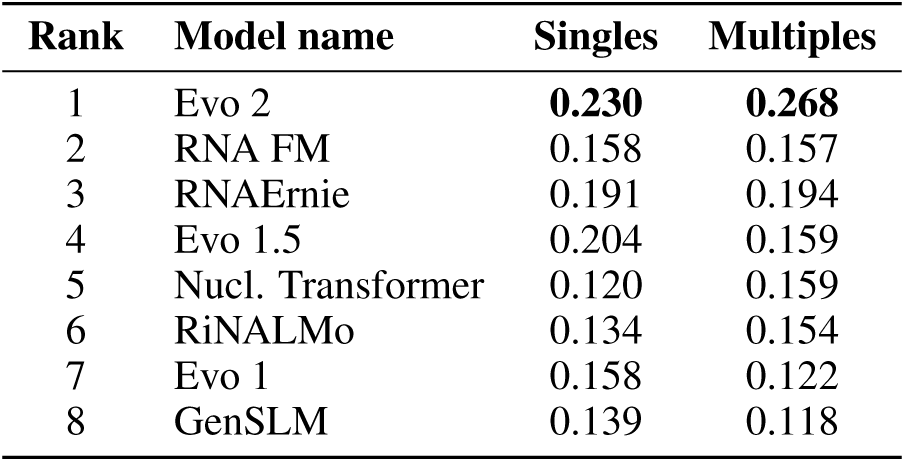
RNAGym - Fitness prediction by mutation type. Average of Spearman’s rank correlation between model scores and experimental measurements by mutation type.

| Rank | Model name | Singles | Multiples |
| --- | --- | --- | --- |
| 1 | Evo 2 | **0.230** | **0.268** |
| 2 | RNA FM | 0.158 | 0.157 |
| 3 | RNAErnie | 0.191 | 0.194 |
| 4 | Evo 1.5 | 0.204 | 0.159 |
| 5 | Nucl. Transformer | 0.120 | 0.159 |
| 6 | RiNALMo | 0.134 | 0.154 |
| 7 | Evo 1 | 0.158 | 0.122 |
| 8 | GenSLM | 0.139 | 0.118 |

**Table A6:** RNA Model Performance (mRNA coding assays) - Spearman Correlation Across Taxa. Spearman correlation between model scores and experimental measurements across different taxonomic groups.

| Rank | Model name | Human | Eukaryote | Prokaryote | Virus | Overall |
| --- | --- | --- | --- | --- | --- | --- |
| 1 | Evo 2 | **0.315** | **0.348** | **0.367** | 0.073 | **0.270** |
| 2 | Evo 1 | 0.095 | 0.112 | 0.323 | 0.079 | 0.161 |
| 3 | Evo 1.5 | 0.114 | 0.128 | 0.318 | 0.042 | 0.157 |
| 4 | RNA-FM | 0.108 | 0.201 | 0.107 | **0.132** | 0.129 |
| 5 | GenSLM | 0.090 | 0.156 | 0.174 | 0.073 | 0.121 |
| 6 | RNAErnie | 0.085 | 0.075 | 0.161 | 0.128 | 0.117 |
| 7 | NT | 0.035 | 0.154 | 0.111 | 0.057 | 0.083 |
| 8 | RiNALMo_score | 0.047 | 0.079 | 0.087 | 0.069 | 0.070 |

**Table A7:** RNA Model Performance (mRNA coding assays) - Area Under Curve (AUC) Across Taxa. AUC metrics for model performance across different taxonomic groups.

| Rank | Model name | Human | Eukaryote | Prokaryote | Virus | Overall |
| --- | --- | --- | --- | --- | --- | --- |
| 1 | Evo 2 | **0.657** | **0.673** | **0.691** | 0.532 | **0.636** |
| 2 | Evo 1 | 0.548 | 0.556 | 0.664 | 0.538 | 0.581 |
| 3 | Evo 1.5 | 0.557 | 0.562 | 0.664 | 0.523 | 0.581 |
| 4 | RNA-FM | 0.558 | 0.603 | 0.555 | 0.562 | 0.566 |
| 5 | GenSLM | 0.548 | 0.582 | 0.594 | 0.539 | 0.565 |
| 6 | RNAErnie | 0.545 | 0.545 | 0.583 | **0.567** | 0.562 |
| 7 | NT | 0.519 | 0.582 | 0.556 | 0.525 | 0.542 |
| 8 | RiNALMo | 0.529 | 0.542 | 0.542 | 0.537 | 0.537 |

**Table A8:** RNA Model Performance (mRNA coding assays) - Matthews Correlation Coefficient (MCC) Across Taxa. MCC metrics for model performance across different taxonomic groups.

| Rank | Model name | Human | Eukaryote | Prokaryote | Virus | Overall |
| --- | --- | --- | --- | --- | --- | --- |
| 1 | Evo 2 | **0.231** | **0.255** | **0.285** | 0.045 | **0.201** |
| 2 | Evo 1.5 | 0.083 | 0.088 | 0.241 | 0.037 | 0.118 |
| 3 | Evo 1 | 0.066 | 0.079 | 0.239 | 0.052 | 0.116 |
| 4 | RNA-FM | 0.090 | 0.152 | 0.080 | 0.091 | 0.098 |
| 5 | RNAErnie | 0.076 | 0.062 | 0.119 | **0.103** | 0.094 |
| 6 | GenSLM | 0.072 | 0.117 | 0.136 | 0.053 | 0.093 |
| 7 | NT | 0.093 | 0.083 | 0.061 | 0.035 | 0.066 |
| 8 | RiNALMo | 0.047 | 0.059 | 0.060 | 0.051 | 0.054 |

**Table A9:** Fitness prediction by RNA type - Difference in Spearman to best score by category. Difference in average of Spearman’s rank correlation between model scores and experimental measurements to the best model by category, by RNA type and overall. The standard error reported corresponds to the non-parametric bootstrap standard error of the difference between the Spearman performance of a given model and that of the best overall model for a given category, computed over 10k bootstrap samples from the set of assays in the RNAGym fitness benchmark.

| Rank | Model | mRNA-splic. |  | mRNA-cod. |  | tRNA |  | Aptamer |  | Ribozyme |  | All |  |
| --- | --- | --- | --- | --- | --- | --- | --- | --- | --- | --- | --- | --- | --- |
|  |  | Diff | SE | Diff | SE | Diff | SE | Diff | SE | Diff | SE | Diff | SE |
| 1 | Evo2 | 0.000 | 0.000 | 0.000 | 0.000 | -0.077 | 0.050 | -0.071 | 0.013 | -0.029 | 0.038 | 0.000 | 0.000 |
| 2 | RNA-FM | -0.328 | 0.126 | -0.141 | 0.034 | 0.000 | 0.000 | 0.000 | 0.000 | 0.000 | 0.000 | -0.058 | 0.028 |
| 3 | RNAErnie | -0.201 | 0.062 | -0.153 | 0.027 | -0.048 | 0.094 | -0.153 | 0.113 | -0.059 | 0.018 | -0.087 | 0.023 |
| 4 | Evo1.5 | -0.300 | 0.125 | -0.113 | 0.031 | -0.079 | 0.050 | -0.146 | 0.089 | -0.032 | 0.034 | -0.099 | 0.021 |
| 5 | NT | -0.311 | 0.014 | -0.188 | 0.036 | -0.147 | 0.043 | -0.044 | 0.022 | -0.054 | 0.027 | -0.113 | 0.025 |
| 6 | RiNALMo | -0.083 | 0.083 | -0.200 | 0.028 | -0.204 | 0.136 | -0.164 | 0.097 | -0.129 | 0.029 | -0.121 | 0.020 |
| 7 | Evo1 | -0.298 | 0.059 | -0.110 | 0.031 | -0.308 | 0.014 | -0.156 | 0.085 | -0.064 | 0.036 | -0.152 | 0.021 |
| 8 | GenSLM | -0.258 | 0.031 | -0.187 | 0.029 | -0.370 | 0.058 | -0.065 | 0.092 | -0.065 | 0.029 | -0.154 | 0.023 |

**Table A10:** Final 2° structure dataset. The secondary structure chemical mapping dataset was further cleaned to only consist of 2A3 and DMS reactivity profiles for RNA in standard condition. The numbers represent the count of chemical mapping profiles while the number in parentheses indicate the count of nucleotides.

| Dataset Type | Train | Test | Total |
| --- | --- | --- | --- |
| 2A3 | 97.6k (11.1M) | 381.3k (43.1M) | 479.0k (54.2M) |
| DMS | 84.6k (9.5M) | 337.6k (37.4M) | 422.2k (46.9M) |
| Total | 182.3k (20.7M) | 718.9k (80.5M) | 901.1k (101.2M) |

**Table A11:**
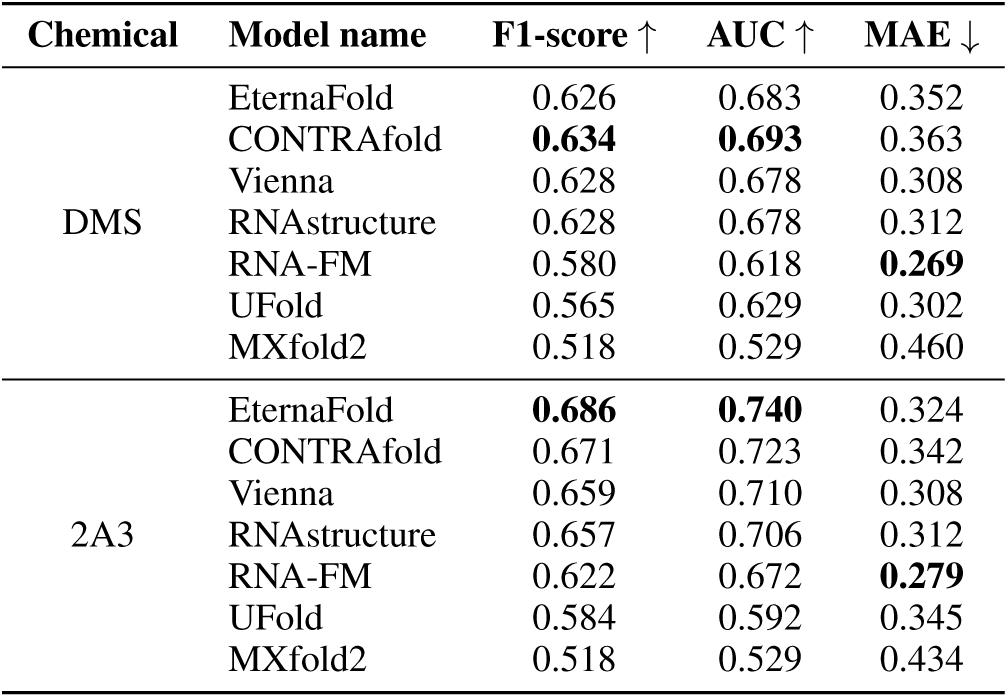
2° structure prediction benchmark performance by chemical mapping type. F1-score, AUC and MAE between model predictions and experimental measurements, by chemical mapping type (DMS and 2A3).

**Table A12:** RNAGym - RibonanzaNet structure prediction benchmark. F1-score, AUC and MAE between RibonanzaNet predictions and experimental measurements (DMS and 2A3) on the RNAGym structure prediction benchmark. From the test dataset split created from RMDB, only sequences from the RibonanzaNet test dataset were kept. The numbers are bolded if they are the best compared to the results in Table 5.

| Chemical | Model name | F1-score ↑ | AUC ↑ | MAE ↓ |
| --- | --- | --- | --- | --- |
| DMS | EternaFold | 0.607 | 0.659 | 0.364 |
|  | CONTRAFold | **0.618** | **0.671** | 0.376 |
|  | Vienna | 0.612 | 0.655 | 0.321 |
|  | RNAstructure | 0.611 | 0.654 | 0.325 |
|  | RNA-FM | 0.560 | 0.589 | **0.284** |
|  | UFold | 0.553 | 0.626 | 0.313 |
|  | MXfold2 | 0.506 | 0.512 | 0.473 |
|  | RibonanzaNet | 0.545 | 0.565 | 0.326 |
| 2A3 | EternaFold | 0.681 | 0.735 | 0.325 |
|  | CONTRAFold | 0.665 | 0.718 | 0.344 |
|  | Vienna | 0.652 | 0.702 | 0.310 |
|  | RNAstructure | 0.649 | 0.698 | 0.315 |
|  | RNA-FM | 0.613 | 0.660 | 0.287 |
|  | UFold | 0.576 | 0.585 | 0.345 |
|  | MXfold2 | 0.517 | 0.530 | 0.438 |
|  | RibonanzaNet | **0.802** | **0.877** | **0.152** |

**Table A13:**
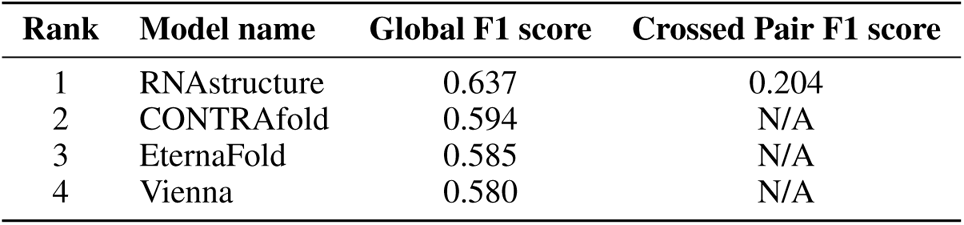
Structure prediction - Pseudoknots. Global and crossed pair F1 score on PseudoBase sequences not related to sequences in the training set (90% sequence identity). 347 sequences were used out of the 358 sequences in the dataset. Crossed pair F1 score is only for the ’crossed’ base pairs (base pairs i-j and m-n with i < m < j < n).

